# Microglia do not restrict SARS-CoV-2 replication following infection of the central nervous system of K18-hACE2 transgenic mice

**DOI:** 10.1101/2021.11.15.468761

**Authors:** Gema M. Olivarria, Yuting Cheng, Susana Furman, Collin Pachow, Lindsay A. Hohsfield, Charlene Smith-Geater, Ricardo Miramontes, Jie Wu, Mara S. Burns, Kate I. Tsourmas, Jennifer Stocksdale, Cynthia Manlapaz, William H. Yong, John Teijaro, Robert Edwards, Kim N. Green, Leslie M. Thompson, Thomas E. Lane

**Affiliations:** Department of Neurobiology & Behavior, School of Biological Sciences, University of California, Irvine 92697; Department of Molecular Biology & Biochemistry, School of Biological Sciences, University of California, Irvine 92697; Department of Psychiatry and Human Behavior, University of California, Irvine School of Medicine, Irvine 92697; Institute for Memory Impairments and Neurological Disorders, School of Biological Sciences, University of California, Irvine 92697; Department of Biological Chemistry, University of California, Irvine School of Medicine, Irvine 92697; Department of Pathology & Laboratory Medicine, University of California, Irvine School of Medicine, Irvine 92697; Department of Immunology & Microbiology, The Scripps Research Institute, La Jolla CA 92037; Center for Virus Research, University of California, Irvine 92697

**Keywords:** SARS-CoV-2, microglia, central nervous system, neuropathology

## Abstract

Unlike SARS-CoV-1 and MERS-CoV, infection with SARS-CoV-2, the viral pathogen responsible for COVID-19, is often associated with neurologic symptoms that range from mild to severe, yet increasing evidence argues the virus does not exhibit extensive neuroinvasive properties. We demonstrate SARS-CoV-2 can infect and replicate in human iPSC-derived neurons and that infection shows limited anti-viral and inflammatory responses but increased activation of EIF2 signaling following infection as determined by RNA sequencing. Intranasal infection of K18 human ACE2 transgenic mice (K18-hACE2) with SARS-CoV-2 resulted in lung pathology associated with viral replication and immune cell infiltration. In addition, ∼50% of infected mice exhibited CNS infection characterized by wide-spread viral replication in neurons accompanied by increased expression of chemokine (*Cxcl9, Cxcl10, Ccl2, Ccl5* and *Ccl19*) and cytokine (*Ifn-λ* and *Tnf-α*) transcripts associated with microgliosis and a neuroinflammatory response consisting primarily of monocytes/macrophages. Microglia depletion via administration of colony-stimulating factor 1 receptor inhibitor, PLX5622, in SARS-CoV-2 infected mice did not affect survival or viral replication but did result in dampened expression of proinflammatory cytokine/chemokine transcripts and a reduction in monocyte/macrophage infiltration. These results argue that microglia are dispensable in terms of controlling SARS-CoV-2 replication in in the K18-hACE2 model but do contribute to an inflammatory response through expression of pro-inflammatory genes. Collectively, these findings contribute to previous work demonstrating the ability of SARS-CoV-2 to infect neurons as well as emphasizing the potential use of the K18-hACE2 model to study immunological and neuropathological aspects related to SARS-CoV-2-induced neurologic disease.

**Importance:** Understanding the immunological mechanisms contributing to both host defense and disease following viral infection of the CNS is of critical importance given the increasing number of viruses that are capable of infecting and replicating within the nervous system. With this in mind, the present study was undertaken to evaluate the role of microglia in aiding in host defense following experimental infection of the central nervous system (CNS) of K18-hACE2 with SARS-CoV-2, the causative agent of COVID-19. Neurologic symptoms that range in severity are common in COVID-19 patients and understanding immune responses that contribute to restricting neurologic disease can provide important insight into better understanding consequences associated with SARS-CoV-2 infection of the CNS.

## Introduction

The clinical spectrum of COVID-19 is complex, and numerous risk factors and comorbidities are considered important in affecting disease severity including age, obesity, chronic respiratory disease, and cardiovascular disease [1]. In addition, neurological symptoms are common in COVID-19 patients, suggesting the virus can potentially infect and replicate in the central nervous system (CNS). Indeed, encephalitis and meningitis have been reported in COVID-19 patients, and viral RNA and protein have been detected within the CSF of infected patients [2-4]. Additionally, human brain organoids are susceptible to SARS-CoV-2 infection [2, 5], yet demonstration of extensive CNS penetrance by SARS-CoV-2 has remained elusive. It is imperative to develop pre-clinical animal models of COVID-19 that capture consistent and reproducible clinical and histologic readouts of many disease-associated symptoms following experimental infection with clinical isolates of SARS-CoV-2 [6]. Importantly, these models should be able to reliably evaluate interventional therapies to limit viral replication and mute immune-mediated pathology, as well as evaluate effectiveness of novel vaccines, all while remaining cost-effective. To date, the most common animal models employed to evaluate COVID-19 pathogenesis include mice, non-human primates (rhesus macaques, cynomolgus macaques and African green monkeys), Syrian hamsters, ferrets, and cats [6].

Human ACE2 (hACE2) transgenic mouse models have provided important insights into the pathogenesis of COVID-19. Perlman and colleagues [7] developed the K18-hACE2 mice, initially used as a mouse model of SARS-CoV-1, which has been successfully employed as a model of COVID-19 [8]. Intranasal inoculation of SARS-CoV-2 in K18-hACE2 mice results in a dose-dependent increase in weight loss and mortality with the lung being the major site of viral infection, while lower amounts of virus are detected in the heart, liver, spleen, kidney, small intestine, and colon [8]. Examination of lungs revealed distribution of viral antigen associated with alveolar damage, interstitial lesions, edema, and inflammation. Lung infection resulted in an increase in expression of interferons as well as inflammatory cytokines and chemokines associated with neutrophil, macrophage/monocyte, and T cell infiltration. Viral RNA was also detected within the sinonasal epithelium, and viral antigen was present in sustentacular cells associated with anosmia [8]. Examination of brains of SARS-CoV-2 infected hACE2 transgenic mice has indicated that infection of the CNS is not consistent, and in some cases, virus is rarely detected [8-12]. This may reflect the SARS-CoV-2 isolate being studied as well as the dose of virus being administered. However, in those animals in which virus penetrates the brain, there can be extensive spread of the virus throughout different anatomic regions accompanied by cell death [8], and these results are consistent with early studies examining SARS-CoV-1 infection of K18-hACE2 mice [7]. High-level of CNS infection in K18-hACE2 is accompanied by meningeal inflammation associated with immune cell infiltration into the brain parenchyma and microglia activation [11]. Enhanced CNS penetrance and replication of SARS-CoV-2 within the CNS of K18-hACE2 is associated with increased mortality; although the mechanisms by which this occurs remain unclear. The present study was undertaken to i) expand on earlier studies examining SARS-CoV-2 infection of human CNS resident cells, ii) evaluate the immune response that occurs in response to SARS-CoV-2 infection of the CNS of K18-hACE2 mice and iii) assess the contributions of microglia in host defense following CNS infection by SARS-CoV-2.

## Results

### SARS-CoV-2 infection of human neurons

Previous studies have indicated neurons are susceptible to infection by SARS-CoV-2 (2); therefore, we infected human iPSC-derived neurons with SARS-CoV-2. Similar to earlier reports, SARS-CoV-2 was able to infect and replicate within neurons as determined by staining for nucleocapsid protein (**Figures 1A and B**). By 48h p.i., viral nucleocapsid protein had spread from the neuron cell body and extended down dendritic and axonal projections (**Figures 1C and D**). Notably, we did not detect syncytia formation in neuron cultures at any time following infection with SARS-CoV-2, suggesting that virus may not spread via fusion with neighboring cells. RNA sequencing analysis revealed that expression of both anti-viral and inflammatory responses in infected neurons was limited relative to the genes within the heatmap at both 24h and 48h post-infection (**Figure 1E**). We then evaluated pathways that may progressively change between 24h and 48h post-infection comparing the Transcripts per million (TPMs) as input for Ingenuity Pathway Analysis (IPA).

**Figure 1.**
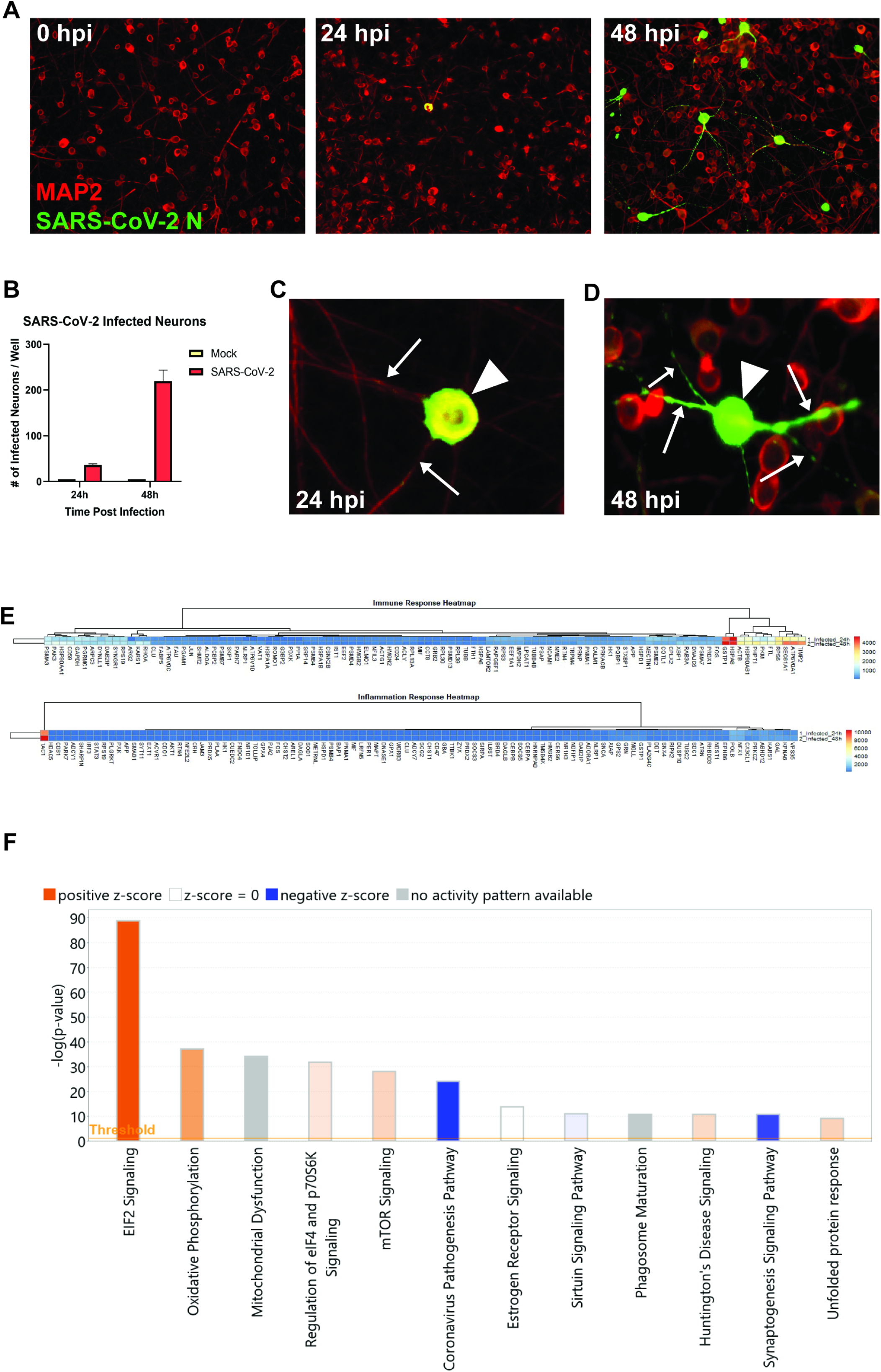
SARS-CoV-2 infects human iPSC-derived neurons. (**A**) hiPSC-derived neurons were infected with SARS-CoV-2 at an MOI of 0.1, immunostained with anti-MAP2 and anti-SARS-CoV-2 N, and imaged at 0, 24, and 48 hours post-infection. (**B**) Quantification of SARS-CoV-2 GFP fluorescence of mock-infected and SARS-CoV-2-infected hiPSC-derived neurons. (**C**) Perinuclear replication of SARS-CoV-2 in neuronal soma (arrowhead) but no viral axonal (arrows) transport at 24 hours post-infection. (**D**) Perinuclear presence of SARS-CoV-2 in soma (arrowhead) and axon (arrows) at 24 hours post-infection. (**E**) Heat map of genes expressed 24 and 48h post-infection. **(F)** Top 12 canonical pathways showing progressive changes from 24 to 48 h post-infection.

**Figure 1F** shows the top 12 IPA canonical pathways that are overrepresented. eIF2 signaling is the predominant pathway induced upon SARS-CoV-2 infection of neurons, followed by pathways associated with oxidative phosphorylation, eIF4, MTOR signaling and mitochondrial dysfunction, the latter with no prediction of activity. Notably, the Coronavirus Pathogenesis Pathway is also overrepresented, however is inhibited in response to neuronal infection by SARS-CoV-2 over the time frame tested with the majority of the genes represented encoding ribosomal proteins (**Figure 1F**).

### SARS-CoV-2 infection of lungs of K18-hACE2 mice

K18-hACE2 mice were intranasally infected with either 1×10^4^, 5×10^4^ or 1×10^5^ plaque-forming units (PFU) of SARS-CoV-2, and weight loss and mortality were recorded. Consistent with other reports [8-12], we observed a general trend towards dose-dependent increase in weight loss and mortality out to day 7 post-infection (p.i.) (**Figure 2A**). qPCR indicated the presence of viral RNA in lungs of infected K18-hACE2 (**Figure 2B**). *In situ* hybridization of lungs of mice infected with 5×10^4^ PFU of virus at day 7 p.i. revealed localized areas of viral infection as determined by expression of spike RNA (**Figures 2C**). Hematoxylin and eosin staining of lungs demonstrated both alveolar and interstitial lesions, with alveolar hemorrhage and edema (**Figures 2D**), interstitial congestion (**Figure 2E**) and lymphocytic infiltrates (**Figure 2F**). qPCR analysis of proinflammatory cytokines and chemokines indicated increased expression of *Ifn-λ, Cxcl10, Cxcr2*, and *Ccl2* yet varied expression of *Il-10, TNF-α, Ccl19, and Ccl5* and limited expression of transcripts for *Ifn-β1*(**Figure 2G**). Hematoxylin and eosin staining showed immune cell infiltrates in SARS-CoV-2-infected lungs containing inflammatory CD8+ T cells as determined by immunofluorescent staining (**Figures 3A-D**). In addition, germinal center-like structures were detected within the lungs of SARS-CoV-2-infected mice with enriched CD8+ T cell accumulation (**Figures 3E and F**). Collectively, our findings are consistent with previous studies employing SARS-CoV-2 infection of K18-hACE2 mice in terms of development of interstitial pneumonia and immune cell infiltration associated with viral RNA present within the lungs [8-12].

**Figure 2.**
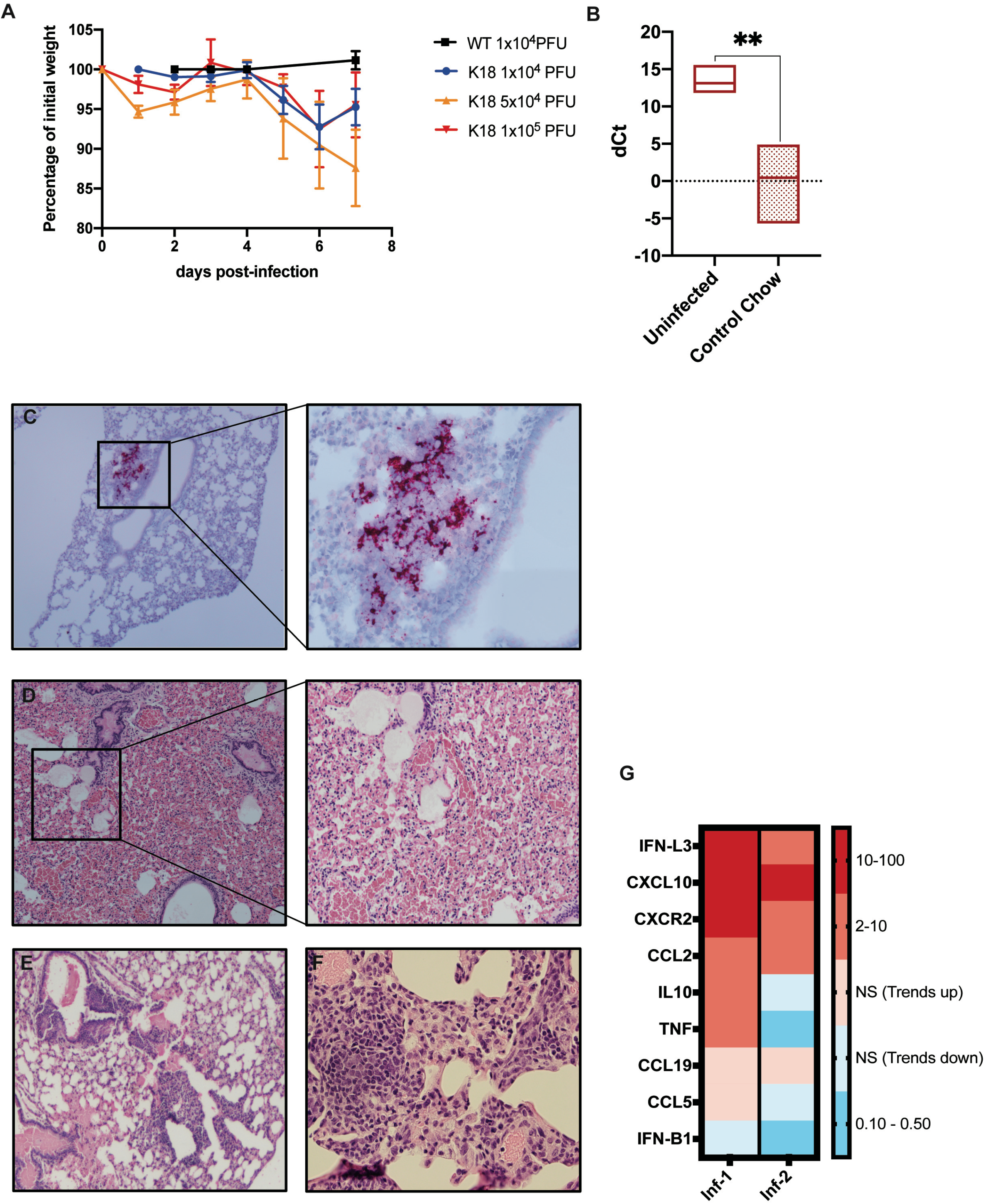
SARS-CoV-2 infection of lungs of K18-hACE2 mice. **(A**) Percent weight change of K18-hACE2 and WT mice infected with indicated dose of SARS-CoV-2. C57BL/6 wildtype (WT) mice (n=3) were infected with 1×104 PFU. K18-hACE2 mice were infected intranasally with SARS-CoV-2 at either 1×10^4^ PFU (n=17), 5×10^4^ PFU (n=4), or 1×10^5^PFU (n=8). (**B**) Quantitative PCR with primers for Spike mRNA on uninfected and SARS-CoV-2-infected mouse lung tissue. dCt values are derived from the difference between the Ct values of Spike mRNA and a housekeeping gene, GAPDH. Lower dCt values indicate increased viral mRNA. (**C**) K18-hACE2 mouse lung tissue at day 7 p.i. with 5×10^4^ PFU SARS-CoV-2 showing localized Spike mRNA expression as determined by RNAscope. Representative H&E images from lungs of SARS-CoV-2-infected mice (5×10^4^ PFU) showing (**D**) airway edema, vascular congestion and intra-alveolar hemorrhage, (**E**) peri-bronchiolar lymphocytic cuffing, and (**F**) Interstitial vascular congestion and lymphocytic infiltrates. (**G**) Quantitative PCR shows the fold changes of the indicated genes in two infected mouse lungs compared to uninfected mice.

**Figure 3.**
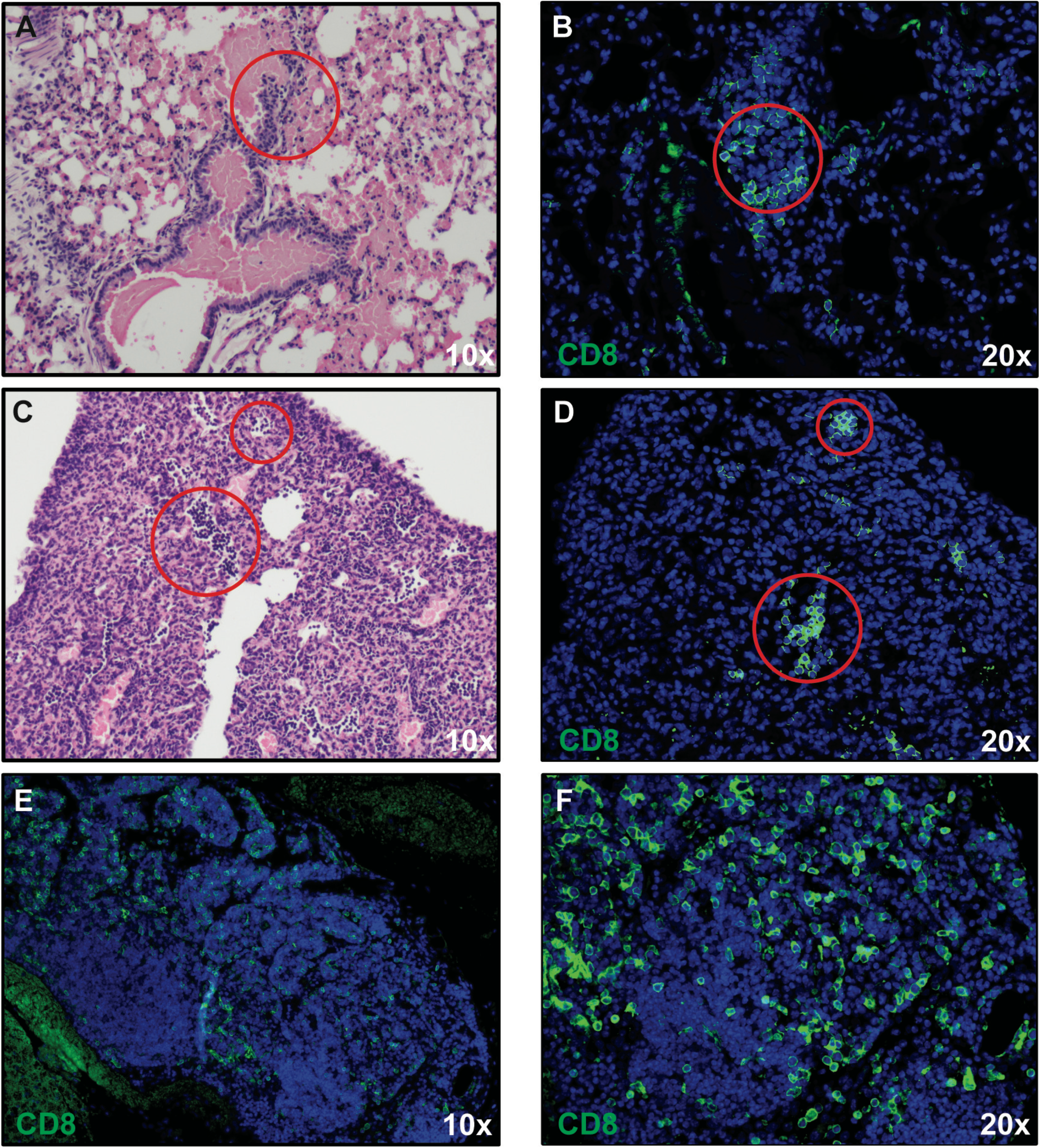
CD8+ T cell infiltration into lungs of SARS-CoV-2-infected mice. H&E staining of lungs of SARS-CoV-2 infected mice at day 7 post-infection reveal inflammation (**A** and **C**) associated with CD8+ T cell infiltration (**B** and **D**) as determined by immunofluorescent staining. Lymph node-like structures were also detected containing CD8+ T cells (**E** and **F**). Panels *A, C*, and *E* 10X magnification; panels *B, D*, and *F* 20X magnification.

### Neuroinvasion by SARS-CoV-2

Following intranasal infection of mice with 5×10^4^ PFU of SARS-CoV-2, virus was detected in the brain as determined by qPCR and RNAscope compared to sham-infected mice (**Figures 4A-C**). Using a Spike-specific probe, we observed wide-spread expansion of viral mRNA throughout the brain with many distinct anatomical regions infected as well as spared. By day 7 p.i., viral RNA was present within the cortex (CTX), striatum (STR), pallidum (PAL), thalamus (TH), hypothalamus (HY), midbrain (MB), pons (P), and medulla (MD), whereas areas that were relatively spared included the olfactory bulb (OB), white matter (WM) tracts, hippocampus (HC) and cerebellum (CB) (**Figure 4B**); no viral RNA was detected in sham-infected K18-hACE2 mice (**Figure 4C**). Demyelinating lesions have been detected in post-mortem brains of COVID-19 patients [13], however we did not detect any evidence of demyelination within the brains of SARS-CoV-2-infected mice as determined by luxol-fast blue (LFB) staining (**Figures 4D and E**). The predominant cellular target for SARS-CoV-2 infection was neurons as determined by cellular morphology of cells positive for viral RNA (**Figures 5A and B**). In a small percentage of infected mice, we were able to detect viral RNA in the olfactory bulb with primary targets being mitral and glomerular neurons (**Figure 5C and D**). Prominent neuropathological changes detected included perivascular cuffing (**Figure 5E**), subventricular inflammation (**Figures 5F**) and leptomeningitis (**Figure 5G**), consistent with previous studies (2, 3).

**Figure 4.**
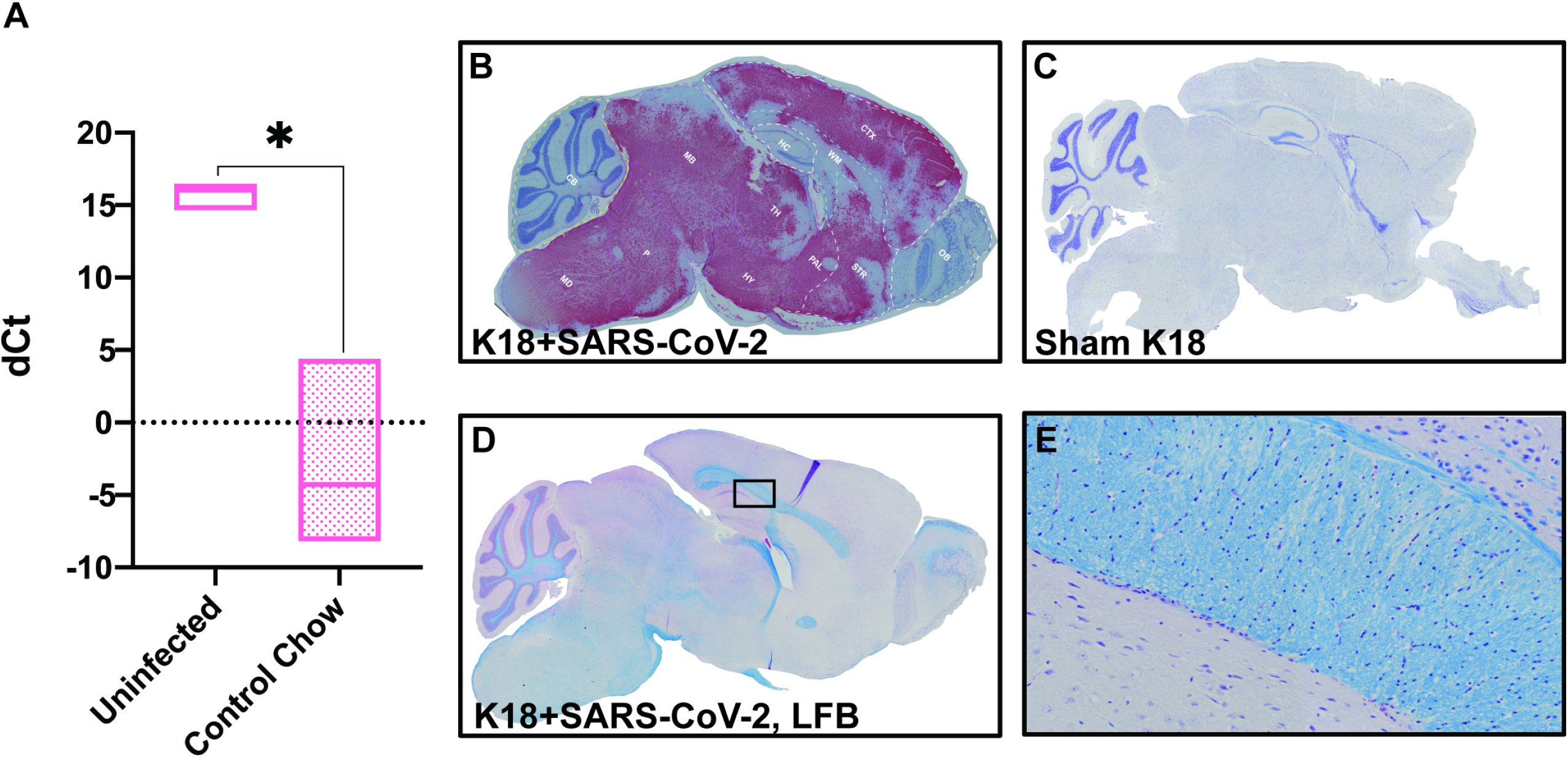
Widespread neuroinvasion by SARS-CoV-2 of K18 human-ACE2 mice. **(A)** Quantitative PCR with primers for Spike mRNA on uninfected and SARS-CoV-2-infected mouse brain tissue at day 7 p.i. dCt values are derived from the difference between the Ct values of Spike mRNA and the housekeeping gene, GAPDH. *In situ* hybridization for Spike viral mRNA in (**B**) SARS-CoV-2-infected and (**C**) sham-infected K18-hACE2 mice. Anatomical regions in which viral RNA is detected (indicated in red) are indicated: cortex (CTX), striatum (STR), pallidum (PAL), thalamus (TH), hypothalamus (HY), midbrain (MB), Pons (P), and medulla (MD), whereas areas that were relatively spared included the olfactory bulb (OB), white matter (WM) tracts and hippocampus (HC) (**D**) Representative brain from SARS-CoV-2 infected brain from panel *B* stained with LFB demonstrates lack of demyelination with (**E**) high-power image of myelin tract showing no inflammation or demyelination.

**Figure 5.**
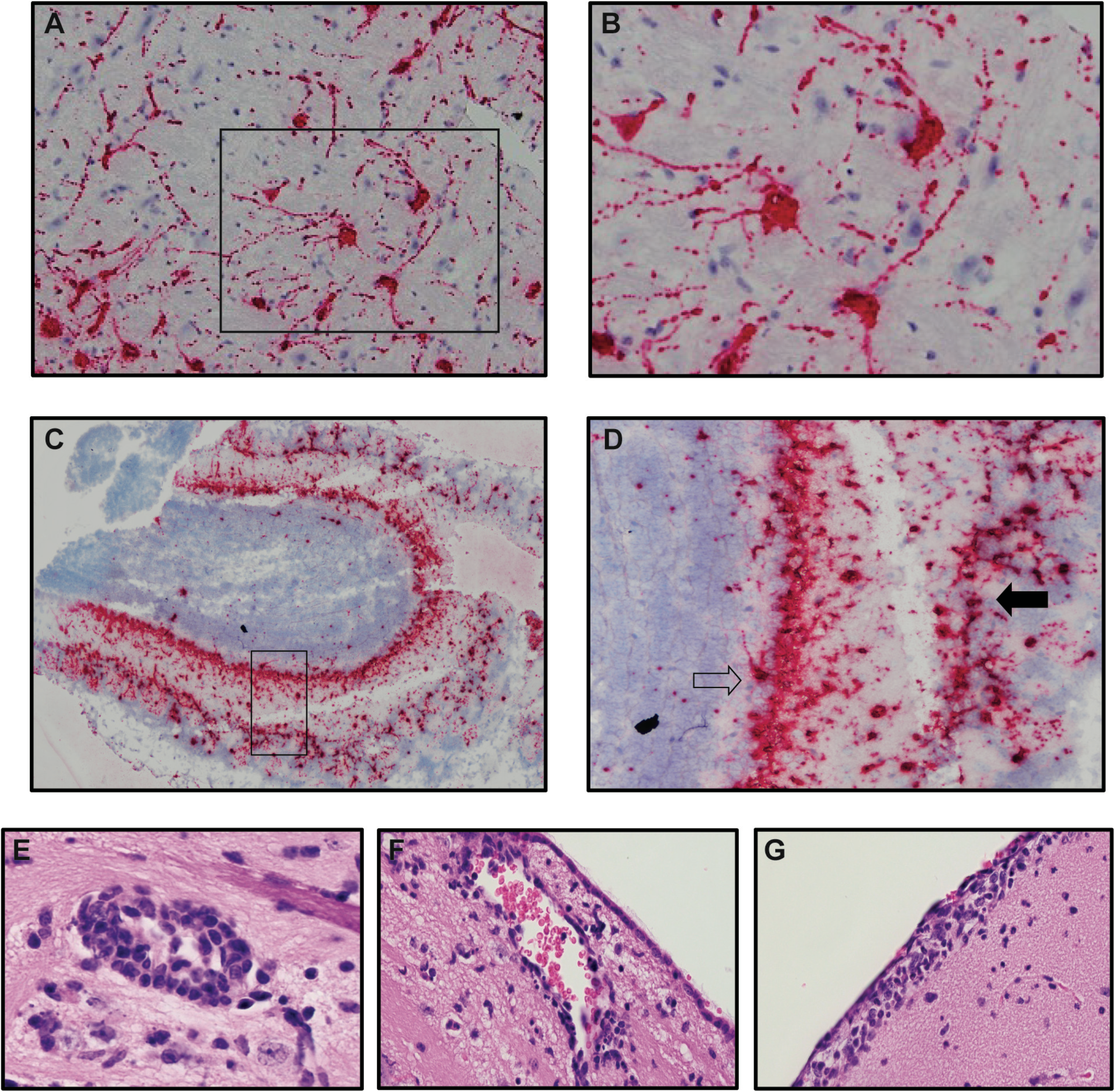
Neurons are targets of infection within the brains of SARS-CoV-2 infected K18-hACE2 mice. Brains of SARS-CoV-2 infected mice at day 7 p.i. were analyzed to assess cellular targets of infection through *in situ* hybridization using RNAscope *in situ* hybridization using Spike-specific probes. (**A**) Cells within the cortex with neuron morphology were primary targets of infection; (**B**) high-power image of cells boxed in panel *A* show viral RNA present within cell body as well as extending down dendrites extending from the cell body. (**C**) Viral RNA was also detected in olfactory bulbs at day 7 p.i. (**D**) high-power image cells boxed in panel *C* reveal neurons in the mitral (open arrow) and glomerular (closed arrow) are infected by virus. Representative H&E images from the brains of infected K18-hACE2 mice at day 7 p.i. depicting (**E**) perivascular cuffing, (**F**) subventricular inflammation, and (**G)** leptomeningitis.

Analysis of CNS myeloid cells showed increased IBA1 soma size and shortened/thickened processes, indicative of microglial activation in SARS-CoV-2-infected mice compared to sham-infected mice (**Figure 6A and B**). In addition, immunofluorescent co-staining for Mac2/galectin-3+ cells, a marker recently identified in peripheral cell infiltrates, and IBA1 (Hohsfield *et al. manuscript in revision*), revealed increased monocyte/macrophage infiltration within the brains of infected mice compared to sham-infected mice (**Figure 6A and B**). Mac2+ cells enter the CNS parenchyma from different anatomic areas including the ventricles, leptomeninges and vasculature (**Figure 6C and D**) when compared to uninfected control mice (**Figure 6E**).

**Figure 6.**
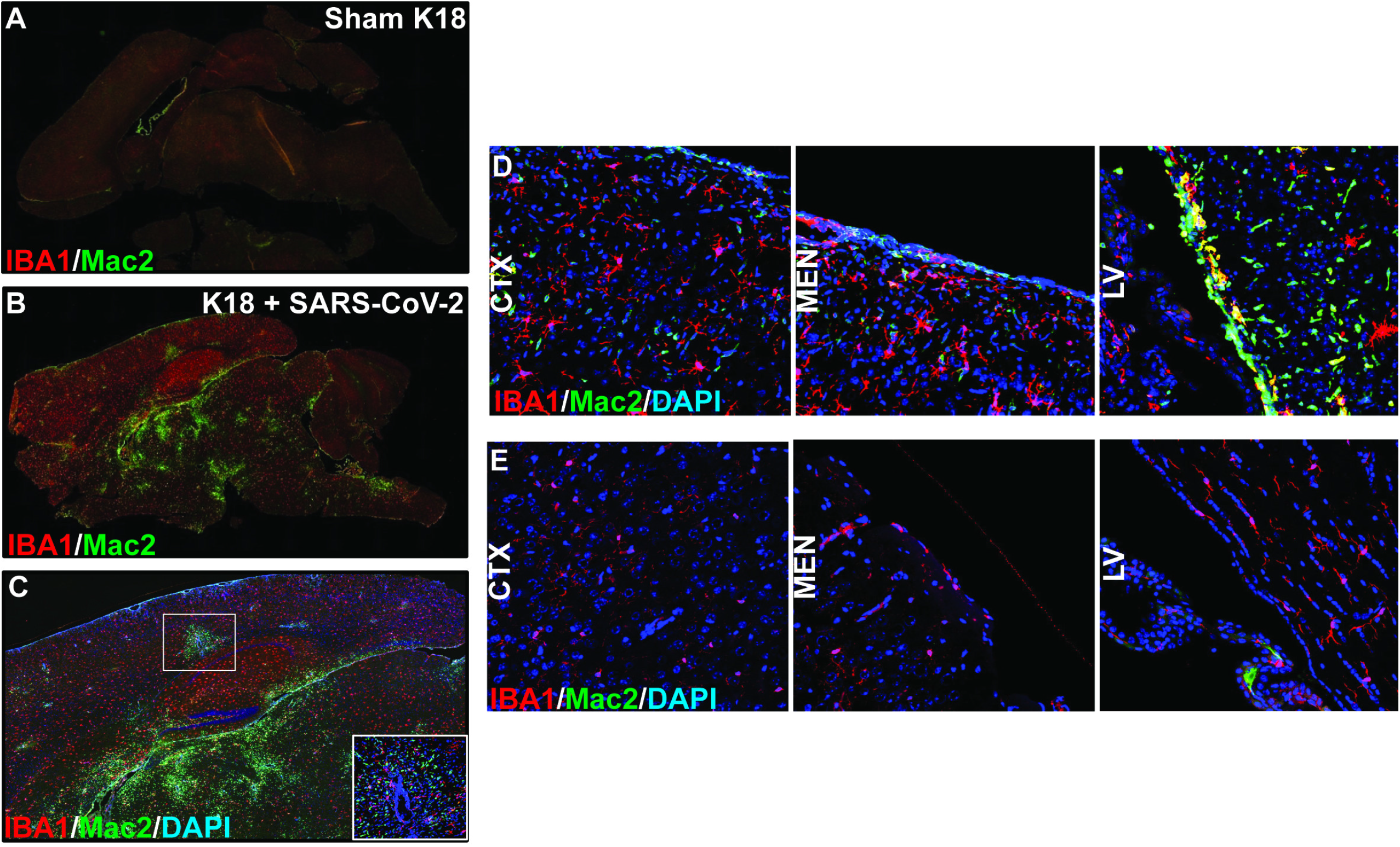
SARS-CoV-2 infection of K18 human-ACE2 mice results in microgliosis and myeloid cell infiltration. Brains from either SARS-CoV-2 or sham-infected mice were removed at day 7 p.i. to evaluate immune cell infiltration. Microglia activation and monocyte infiltration were determined in sham-infected mice **(A)** and SARS-CoV2 infected mice **(B)** by staining for expression of Iba1 (red) and Mac2/galectin 3 (green), respectively. Infiltration of peripheral monocytes into the SARS-CoV-2 infected brain parenchyma occurs via the vasculature (**C**) as well as ventricular and leptomeningeal spaces (**D**) compared to uninfected control mice (**E**).

### Microglia ablation does not affect disease or control of viral replication in the CNS

Previous studies have implicated the importance of microglia in aiding in control of neuroadapted murine beta-coronaviruses by enhancing anti-viral T cell responses through augmenting antigen presentation (11-13). Notably, microglia are dependent on signaling through the colony stimulating factor 1 receptor (CSF1R) for their survival and can be effectively eliminated with CSF1R inhibitors that cross the blood brain barrier [14]. K18-hACE2 mice were fed pre-formulated chow containing either the CSF1R inhibitor PLX5622 (1,200 ppm) [15] or control chow 7 days prior to intranasal infection with 5×10^4^ PFUs of SARS-CoV-2, and remained on drug for the duration of the experiment [6]. No notable differences were detected in weight loss between experimental groups (**Figure 7A**). SARS-CoV-2-infected mice treated with either PLX5622 or control chow were sacrificed at day 7 p.i., and viral mRNA levels in lungs and brains were determined. As shown in **Figure 7B**, we detected no significant differences in the Spike mRNA transcripts in either the lung or the brain between experimental groups.

**Figure 7.**
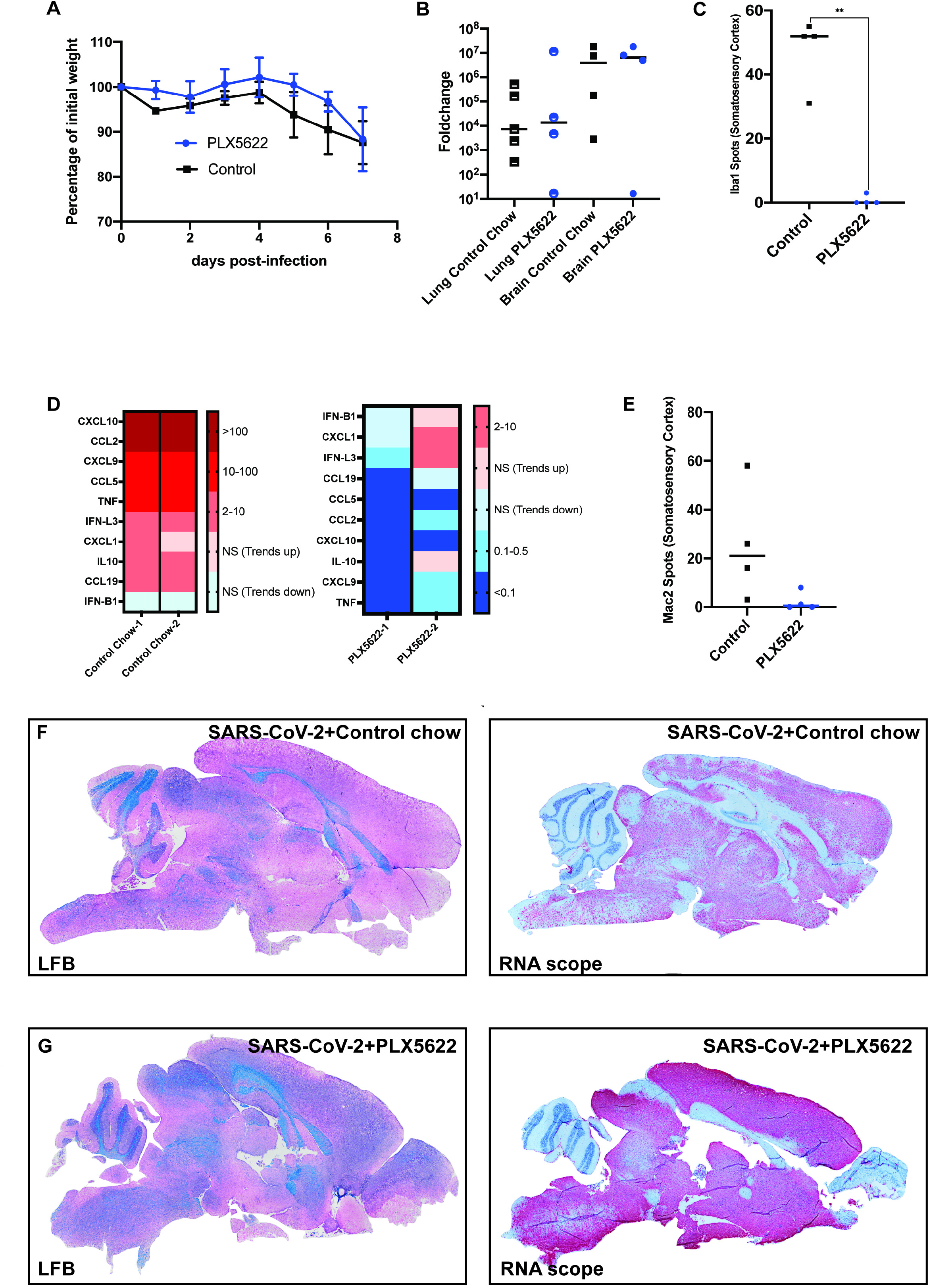
Microglia ablation does not impact control of viral replication in the CNS. (**A**) Weight loss of K18-hACE2 mice infected intranasally with 5×10^4^ PFU of SARS-CoV-2 that were fed either control chow (n=4) of PLX5622-formulated chow (n=4). (**B**) Quantitative PCR in the brains and lungs of infected PLX-treated and non-PLX-treated mice shows no significant difference in the levels of Spike mRNA in either lung or brain tissue as a result of PLX5622 treatment. **(C)** Quantification of Iba1-positive cells in the somatosensory cortex shows a significant (**p<0.01) depletion of microglia from PLX5622 treated mice compared to control mice. (**D**) qPCR analysis of brains of experimental mice indicated a reduction in expression of pro-inflammatory cytokines/chemokines in the brains of PLX5622-treated mice compared to control mice. **(E)** Quantification of Mac2-positive cells in the somatosensory cortex an overall trend in reduced numbers in PLX5622-treated mice compared to controls. Brains from SARS-CoV-2-infected mice treated with either (**F**) control chow or (**G**) PLX5622 were stained with LFB to assess demyelination or the presence of viral RNA determined by RNAscope. Representative brain sections are from experimental mice at day 7 p.i.

Quantification of Iba1-positive cells revealed a >95% reduction (p<0.01) in microglia in PLX5622-treated mice compared to control mice. Overall, PLX5622-mediated ablation of microglia resulted in a dramatic reduction in expression of proinflammatory cytokine/chemokine genes within the brains of SARS-CoV-2 infected mice when compared to infected mice fed control chow (**Figure 7D**). By day 7 p.i., SARS-CoV-2 infection resulted in increased expression of proinflammatory cytokines and chemokines in mice treated with control chow. The highest transcript levels were for the T cell chemoattractants CXCL9 and CXCL10; monocyte/macrophage chemoattractants, CCL2 and CCL5; and TNF-α (**Figure 7D**). Transcripts encoding the neutrophil chemoattractant CXCL1, and B cell chemoattractant CCL19 were also increased, as well as IFN-λ type 3 and IL-10 when compared to uninfected controls (**Figure 7D**).

Depletion of microglia resulted in a marked reduced expression of the majority chemokine/cytokines transcripts. While not significant, there was a marked reduction in Mac2/galactin3+ cells in PLX5622-treated mice when compared to control chow and this correlated with reduced expression of monocyte/macrophage chemoattractant chemokines CCL2 and CCL5 (**Figure 7E**) that have previously been shown to attract these cells into the CNS of mice infected with a neurotropic mouse coronavirus [16-21].

## Discussion

In the face of the ongoing COVID-19 pandemic, it is imperative to develop pre-clinical animal models of COVID-19 that capture consistent and reproducible clinical and histologic readouts of many disease-associated symptoms following experimental infection with SARS-CoV-2 [1]. For both SARS-CoV-2 and SARS-CoV-1, the surface-bound viral spike glycoprotein uses the cellular surface receptor protein, ACE2, to bind and enter cells. However, mouse ACE2 does not efficiently bind the spike glycoprotein of either SARS-CoV-1 or SARS-CoV-2, rendering wildtype mice not useful in the study of SARS or COVID-19 pathogenesis, respectively, due to an inefficient ability to infect and replicate in host cells. Human ACE2 (hACE2) transgenic mouse models have provided important insights into the pathogenesis of COVID-19 in terms of evaluating the efficacy and duration of immune responses elicited in response to infection as well as testing vaccines, anti-viral drugs and monoclonal antibody therapies to restrict viral replication and limit disease severity [8].

Intranasal inoculation of K18-hACE2 mice with SARS-CoV-2 resulted in weight loss along with viral infection and replication within the lungs that was associated with a robust inflammatory response. These findings are consistent with other reports that demonstrated the presence of neutrophils, monocytes/macrophages and T cells within the lungs of SARS-CoV-2-infected K18-hACE2 mice [8, 11]. In response to infection, expression of proinflammatory cytokines/chemokines was increased and correlated with the presence of inflammatory cells. We detected inflammatory CD8+ T cells in the lungs of infected mice and these cells are presumably responding to the T cell chemoattractant CXCL10. Our lab has previously shown that induction of CXCL10 expression following experimental infection of mice with a neuroadapted strain of murine coronavirus is critical in host defense and acts by attracting virus-specific T cells into the

CNS [22-24]. Similarly, inflammatory monocyte/macrophages are likely recruited in response to expression of CCL2 as this has been shown to attract these cells following murine coronavirus infection of mice [16-19]. The increased expression of transcripts encoding CXCR2 most likely reflects the presence of inflammatory neutrophils [25]. Therapeutic targeting of these chemokines may therefore alter immune cell trafficking into the lungs of infected mice and alleviate the severity of lung pathology. How blocking chemokine signaling would affect SARS-CoV-2-induced lung pathology in COVID-19 patients is currently under investigation.

Clinical reports of COVID-19 patients cite a dysregulated immune response characterized by elevated chemokine expression as key in the development of pathogenesis [26, 27]. Similar to our findings using the K18-hACE2 model, these reports have found upregulation of CCL2 (MCP1), CXCL8 (IL-8), and CXCL10 (IP10) to correlate with severity of disease and have suggested evaluation of these as biomarkers for disease and as targets for therapeutic intervention [28-30]. Leronlimab, a CCR5-blocking antibody, is currently in phase 2 clinical trials in the U.S. for treatment of mild, moderate, and severe COVID-19 (NCT04343651, NCT04347239). In addition, limiting neutrophil infiltration into the lungs of COVID-19 patients may help prevent disease progression and outcome. With this in mind, the monoclonal antibody BMS-986253 that blocks CXCL8 as well as Reparixin, an oral inhibitor of CXCR1/2, are both currently undergoing clinical trials in the U.S. for reducing the severity of COVID-19-related pneumonia by reducing neutrophil accumulation within the lungs of infected patients (NCT04878055, NCT04347226). A clinical trial was recently completed using measurement of CXCL10 in a Clinical Decision Support Protocol in COVID-19 patients (NCT04389645) that positively correlated CXCL10 levels with mortality suggesting that targeting CXCL10 signaling may also be beneficial in managing disease severity [31].

Examination of brains of SARS-CoV-2-infected K18-hACE2 mice, as well as brains from other hACE2 transgenic mice, has revealed that virus is able to infect, replicate and spread within the parenchyma and this is considered important in contributing to mortality [2, 5, 8, 9]. Virus can be detected in different anatomic regions of the brain and is accompanied by cell death [5] and these observations are consistent with early studies examining SARS-CoV-1 infection of K18-hACE2 mice [7]. Our results indicate that neurons are primary targets for SARS-CoV-2 infection within the brains of K18-hACE2 mice which is consistent with other studies [10, 32, 33]. Furthermore, human iPSC-derived neurons were susceptible to SARS-CoV-2 infection and virus was able to replicate in these cells. The detection of viral antigen within dendrites extending from the cell body suggests this may offer a unique mechanism of viral replication and spread in neurons and this is further emphasized by the absence of cell death and/or syncytia formation. RNA sequencing of infected neurons revealed that expression of most inflammatory and anti-viral response genes were reduced in SARS-CoV-2-infected neurons, consistent with a previously published study highlighting the muted immune response in neurons infected with SARS-CoV-2 as compared to Zika virus [2]. However, IPA analysis determined that despite the lack of induced immune and inflammatory response in infected neurons, there was a significant overrepresentation of pathways associated with eIF2, oxidative phosphorylation, regulation of eIF4 and mTOR signaling. eIF2, which mediates initiation of eukaryotic translation by binding Met-tRNA_i_^Met^ to the ribosomal subunit, can be activated via phosphorylation by various kinases, including protein kinase-RNA-dependent (PKR), PKR-like endoplasmic reticulum kinase (PERK), and heme regulated inhibitor (HRI) [34]. Upregulation of eIF2 phosphorylation by these can occur in response to various types of stressors including viral infection, ER stress, or oxidative stress, respectively [34]. Upregulation of eIF4, which mediates recruitment of ribosomes to mRNA for translation, alongside overactivation of mTOR signaling, which modulates eIF4 activity, suggests infection of neurons with SARS-CoV-2 induces increased cellular activity and protein production even in the absence of inflammatory and anti-viral gene induction [35]. Whether this increase in protein production pathways corresponds to viral hijacking of cellular machinery for viral reproduction or if it corresponds to other induced mechanisms of neuronal stress/survival response remains to be determined.

Resident cells of the CNS are important in host defense following viral infection through either secretion of anti-viral cytokines like type I interferon (IFN-I) and pro-inflammatory cytokines/chemokines, and presentation of antigen to infiltration of antigen-sensitized T cells. We found the transcripts encoding the T cell chemoattractant chemokines including CXCL9 and CXCL10 were expressed along with monocyte/macrophage chemokines CCL2 and CCL5 and CXCL1 which attracts neutrophils. In addition, transcripts specific for the B cell chemoattractant CCL19 were also detected. The chemokine elicited in response to CNS infection of K18-hACE2 by SARS-CoV-2 infection is remarkably similar to the chemokine response following CNS infection of mice with the neuroadapted strains of murine coronaviruses [36, 37]. This is interesting in that murine coronavirus are primarily tropic for glial cells e.g. astrocytes, microglia and oligodendrocytes with relative sparing of neurons while neurons appear to be exclusively targeted by SARS-CoV-2. These findings argue that expression of chemokines following coronavirus infection of the CNS may not be influenced by the cellular target of infection and this may reflect a localized response to expression of interferons (IFN) expressed in response to viral infection. With this in mind, we did detect IFNλ3 transcripts within the brains of SARS-CoV-2 infected mice yet IFNβ1 transcripts was noticeably reduced. The inflammatory response consisted primarily of monocytes/macrophages as determined by immunofluorescent staining for Mac2 and these cells are most likely migrating into the CNS in response to CCL2 and CCL5 expression [18-21]. We did not detect a robust T cell response and this was surprising given the expression levels of CXCL9 and CXCL10 transcripts. Whether this simply reflected that T cells had yet to migrate into the brains of infected mice at the time of sacrifice or if efficient translation of these transcripts is compromised is not known at this time.

Microglia are now recognized to be important in host defense in response to viral infection of the CNS [38-42]. Targeted depletion of microglia via CSF1R inhibition leads to increased mortality in mice infected with West Nile Virus (WNV) and is associated with diminished activation of antigen presenting cells (APCs) and limited reactivation of virus-specific T cells that leads to reduced viral clearance [39, 41]. Similar findings have been reported for other neurotropic viruses including Japanese encephalitis virus (JEV) [41] and Theiler’s murine encephalomyelitis virus (TMEV) [40, 42]. Moreover, microglia have also been shown to enhance host defense following CNS infection by the neurotropic JHM strain of mouse hepatitis virus (JHMV, a murine coronavirus) and this was related to inefficient T cell-mediated control of viral replication [38, 43]. Additionally, the absence of microglia also results in an increase in the severity of demyelination, accompanied by a decrease in remyelination [43, 44]. In response to SARS-CoV-2 CNS infection, we did detect microgliosis indicating these cells are responding to infection and may be involved in host defense. Ablation of microglia via PLX5622 administration did not affect clinical disease nor viral burden within the brain indicating these cells are dispensable in terms of controlling SARS-CoV-2 replication within the brain in the K18-hACE2 model. There was a marked reduction in expression of proinflammatory cytokines/chemokines including monocyte/macrophage chemoattractants CCL2 and CCL5 and this corresponded with an overall reduction in numbers of these cells in the brains of PLX5622-treated mice. These findings support the notion that microglia do contribute to the neuroinflammatory response, in part, through influencing expression of chemokines/cytokines in response to SARS-CoV-2 infection of the CNS of K18-hACE2 mice.

Very early in the COVID-19 pandemic, it became apparent that an abundant number of patients exhibited a variety of neurologic conditions that ranged in severity. Although numerous neurological symptoms have been associated with COVID-19, the most common neurologic symptoms include anosmia/dysgeusia, delirium, encephalopathy, and stroke [45-47]. The overwhelming evidence indicates that SARS-CoV-2 is not readily detected within the CNS by either PCR and/or immunohistochemical staining [2, 48] suggesting that neurologic complications associated with COVID-19 patients may occur through alternative mechanisms, including the potential development of autoreactive antibodies specific for neural antigens [49]. COVID-19 patients have anti-SARS-CoV-2 IgG antibodies in the cerebral spinal fluid (CSF) that recognized target epitopes that different from serum antibodies. Moreover, a portion of COVID-19 patients exhibited CSF antibodies that targeted self-antigens, arguing for the possibility that neurologic disease may be associated with CNS autoimmunity [49]. Nonetheless, neurons and cells of the vasculature have been shown to be targets of infection [50-54]. Despite increasing evidence indicating extensive SARS-CoV-2 infection of the CNS does not occur, autopsy findings indicate the presence of microglia nodules, astrocyte activation and CD8+ T cell infiltration in the brain, providing evidence for immune responses occurring within the CNS of infected patients [3, 55, 56]. Related to this, a recent study indicated microglia nodules interacting with inflammatory CD8+ T cells within distinct anatomical regions of the brains of COVID-19 patients and this correlated with alerted systemic inflammation [57]. It is incontrovertible that transgenic hACE2 models, particularly the K18-hACE2 model, have provided a better understanding of the pathogenesis of COVID-19 yet the one notable and consistent drawback of many of these models is the ability of virus to infect, efficiently replicate, and spread within the parenchyma, which contributes to increased mortality. These findings emphasize the importance of working with animal models in which SARS-CoV-2 entry into the CNS is more consistent with what has been observed in COVID-19 patients.

## Materials and Methods

### Mice and viral infection

All experiments were performed in accordance with animal protocols approved by the University of California, Irvine Institutional Animal Care and Use Committee. 8-16 week-old heterozygous K18-hACE2 C57BL/6 [strain: B6.Cg-Tg(K18-ACE2)2Prlmn/J] mice were obtained from Jackson Laboratory. Animals were housed by sex in single use disposable plastic cages and provided ad-libitum water. SARS-CoV-2 isolate USA-WA1/2020 was obtained from BEI. Mice were inoculated with between 10^4^-10^5^ PFU of SARS-CoV-2 in 10µL of DMEM or sham inoculated. Inoculations were performed under deep anesthesia through an intraperitoneal injection of a mixture of ketamine and xylazine. Infected and uninfected mice were examined and weighed daily. Animals were euthanized early if they reached pre-determined euthanasia criteria.

### iPSC-neuronal differentiation

iPSC line iCS83iCTR33n1 was previously derived, characterized [58] and maintained as described at 37°C, 5% CO_2 -_ on hESC Matrigel^®^ (Fisher Scientific cat#08774552) with daily feeding of mTeSR1™ (Stem Cell Technologies cat#85850) [59]. Neuronal differentiation was performed using the protocol as described [59] with a modification that the cells were frozen at the neural progenitor stage at day 8 in CryoStor CS10 (Stem Cell Technologies #07931). Cells were thawed into LIA medium (ADF supplemented with 2mM Glutamax, 2% B27 without vitamin A, 0.2µM LDN 193189 and 1.5µM IWR1 20ng/mL Activin A (Peprotech #120-14E)) containing 10µM Y-27632 dihydrochloride for the first day post-thaw for further neural differentiation and subsequent daily feeds with LIA without Y-27632 dihydrochloride. Differentiated neurons were plated at 1×10^6^ cells per well in 6-well plate format or at 8×10^4^cells per chamberslide well for imaging. Cells were infected with either pseudovirus or SARS-CoV-2 at d46-50 after start of differentiation.

### SARS-CoV-2 infection of iPSC-derived neurons

SARS-CoV-2 isolate USA-WA1/2020 was obtained from BEI. Media was removed from cells and replaced with virus-containing media (or non-virus media for mock-infection wells) at 500µL per well of a 6-well plate or 200µL per chamberslide well for 1hr adsorption at 37°C, 5% CO_2_, for infection of cells at MOI = 0.1. Culture plates and chamber slides were gently rocked every 15min to ensure even distribution of infection media. Following 1hr adsorption, non-virus containing media was added to all cells for final volume of 2mL per well of a 6-well plate or 700µL per chamberslide well and were allowed to incubate with virus for 24 or 48h and then fixed with 4% PFA for subsequent immunocytochemical staining. Cells in 6-well plates were allowed to incubate with virus for 24, 48, or 72h; supernatants were then collected, and cells harvested using 700µL of cold TRIzol Reagent (Ambion, 15596018) for subsequent qPCR analysis.

### PLX5622 treatment

Rodent chow (AIN-76A) formulated with CSF1R inhibitor-PLX5622 at a dose of 1,200 ppm was provided by Plexxikon, Inc (Berkeley, CA). Mice were fed either PLX5622 chow or control chow 7 days prior to viral infection, and chow was continued until mice were sacrificed at defined time points post-infection.

### RNA extraction

All RNA from VSV experiments with iPSC neurons, astrocytes, and microglia was extracted via the RNeasy Mini Kit (Qiagen, 74106) using the “Purification of Total RNA from Animal Cells using Spin Technology” protocol. Homogenization was performed using QIAshredder spin columns (Qiagen, 79656). RNA from SARS-CoV-2 infected neurons was extracted via the RNeasy Mini Kit using the “Purification of Total RNA, Including Small RNAs, from Animal Cells” protocol. TRIzol was substituted for QIAzol, and Buffer RW1 was substituted for Buffer RWT. Homogenization was performed using QIAshredder spin columns. All RNA from mouse tissues was extracted via the RNeasy Mini Kit using the “Purification of Total RNA, Including Small RNAs, from Animal Tissues” protocol. TRIzol was substituted for QIAzol, and Buffer RW1 was substituted for Buffer RWT. Homogenization was performed using the Bead Ruptor 12 (Omni International) and 1.4 mm ceramic beads (Omni International, 19-627). For brain tissue, the machine was set to 2 cycles at 2.25m/s for 15 seconds with a 1 second pause between cycles. For lungs, the machine was set to 2 cycles at 2.4m/s for 20 seconds with a 1 second pause between cycles.

### cDNA synthesis

All cDNA was made by following the “First Strand cDNA Synthesis” standard protocol provided by New England Biolabs with their AMV Reverse Transcriptase (New England Biolabs, M0277L). Random hexamers (Invitrogen, N8080127) were used for the reactions. RNase inhibitors were not used in the cDNA synthesis.

### Gene expression analysis via quantitative PCR

All qPCRs were performed using the Bio-Rad iQ5 and iTaq™ Universal SYBR® Green Supermix (Bio-Rad, 1725120). The standard protocol by Bio-Rad for iTaq™ Universal SYBR® Green Supermix was used unless otherwise stated. Reactions were 10µL, and the machine was set to run for 1 cycle (95°C for 3 minutes), followed by 40 cycles (95°C for 10 seconds, then 55°C for 30 seconds). The following primer sequences were used:

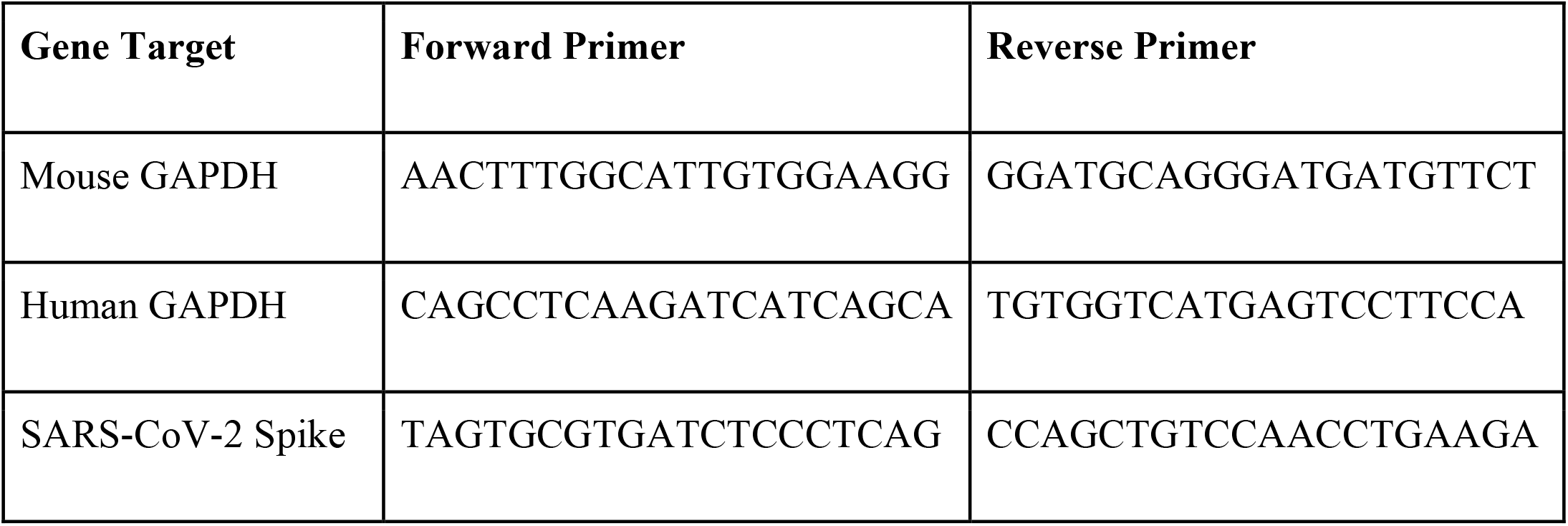

For the brain cytokine and chemokine qPCR, Qiagen’s custom qPCR arrays were employed, following the protocol, “Real-Time PCR for RT^2^ Profiler PCR Arrays Formats A, C, D, E, F, G.” The plates were pre-aliquoted with primers for the following murine genes: Glyceraldehyde-3-phosphate dehydrogenase (GAPDH), beta actin, chemokine ligand 10 (CXCL10), chemokine ligand 9 (CXCL9), chemokine ligand 2 (CCL2), chemokine ligand 5 (CCL5), interferon gamma (IFN-γ), interferon beta-1 (IFN-B1), tumor necrosis factor (TNF-α), interleukin 10 (IL10), chemokine ligand 1 (CXCL1), interleukin 28B (IFN-L3), chemokine ligand 19 (CCL19). Reactions were 25µL (1µL cDNA, 11.5 µL UltraPure Distilled Water (Invitrogen, 10977-015), 12.5 µL iTaq™ Universal SYBR® Green Supermix). The machine was set to run for 1 cycle (95°C for 10 minutes), followed by 40 cycles (95°C for 15 seconds, then 60°C for 1 minute). Ct values for each sample were normalized to an internal control (GAPDH), yielding the dCt values. dCt values of infected or PLX-treated samples were compared to appropriate control samples, as indicated, to produce ddCt values. The relative fold change between samples used in the ddCt calculation was calculated (2^-ddCt^).

### qPCR statistical analysis

Statistical analysis was performed in Prism (GraphPad Software). The Brown-Forsythe and Welch’s ANOVA tests were used to compare the means between groups. Dunnett’s test to correct for multiple comparisons was performed when necessary.

### mRNA-Seq

Total RNA was isolated using the Qiagen RNeasy Kit and QIAshredders for cell lysis. One microgram of RNA with RNA integrity number values >9 was used for library preparation using the strand-specific Illumina TruSeq mRNA protocol. Libraries were sequenced on the NovaSeq 6000 platform using 100 cycles to obtain paired-end 100 reads at >30 million reads per sample. For RNA-seq analysis, fastq files were trimmed using a base quality score threshold of >20 and aligned to the hg38 genome with Hisat 2. Reads passing quality control were used for quantification using featureCounts and analyzed with the R package DESeq2 to identify DEGs. Genes passing an FDR of 10% were used for GO enrichment analysis using GOrilla (27) (http://cbl-gorilla.cs.technion.ac.il/). For the heatmaps, a list of genes from GO:0006955 immune response and GO:0006954 inflammatory response was used. From those lists, the genes with the top 100 most variable TPM were plotted using the R package pheatmap. For the 48h vs 24h IPA, the TPM of all genes in the 1_Infected_24h sample were subtracted from the TPM of all genes in the 2_Infected_48h sample. The list of genes were sorted by the difference and the top 400 and bottom 400 selected to use as input for IPA’s core analysis.

### Histology and immunohistochemical staining

Mice were euthanized at defined time points post-infection and tissues harvested according to IACUC-approved guidelines. Tissues were collected and placed in either cold TRIzol for qPCR analysis or 4% PFA for fixation and subsequent histological analysis. Following fixation, the 24-48hr brains were either cryoprotected in 30% sucrose, embedded in O.C.T. (Fisher HealthCare), and sliced via Cryostat in 10µm sagittal sections. Alternatively, tissues were dehydrated and paraffin-embedded and subsequently use for RNAscope or hematoxylin/eosin (H&E) in combination with luxol fast blue (LFB) to assess demyelination within the brains of experimental mice. For immunohistochemical staining of O.C.T.-embedded tissues, slides were rinsed with PBS to remove residual O.C.T., and antigen retrieval (incubation with 10mM sodium Citrate at 95°C for 15min) was performed if required for specific antigens at which point samples were incubated with 5% normal goat serum and 0.1% Triton-X, followed by overnight incubation at 4 °C with primary antibodies. Several primary antibodies were used, including Iba1 (1:500 Wako), GFAP (1:1000 Abcam), Mac2/ Galactin-3 (1:500, CL8942AP Cedarlane), CD4 (1:200 Abcam), CD8 (1:200 Abcam), MHC I (1:200 Abcam), and MHC II (1:200 Abcam). On the second day, slides were treated with appropriate secondary antibodies (1:1000 goat anti-rat/rabbit Invitrogen, 1:1000 goat anti-chicken, Abcam) following PBS rinsing. Slides were then mounted with DAPI Fluoromount-G (SouthernBiotech). High-resolution fluorescent images were obtained using a Leica TCS SPE-II confocal microscope and LAS-X software. For whole brain stitches, automated slide scanning was performed using a Zeiss AxioScan.Z1 equipped with a Colibri camera and Zen AxioScan 2.3 software. Microglial morphology was determined using the filaments module in Bitplane Imaris 7.5, as described previously (Elmore, Lee, West, & Green, 2015). Cell quantities were determined using the spots module in Imaris.

### RNAscope in situ hybridization of SARS-CoV-2 spike RNA

RNA in situ hybridization was performed via RNAscope 2.5 HD Red Assay Kit (Advanced Cell Diagnostics, Cat: 322350) in accordance with manufacturer’s instructions. Fixed tissue sections were treated with the manufacturer’s Fresh Frozen Tissue Sample Preparation Protocol, fixed in chilled 4% PFA, dehydrated, and treated with H_2_O_2_ and Protease IV before probe hybridization. Paraffinized sections were deparaffinized and treated with H_2_O_2_ and Protease Plus prior to hybridization. Probes targeting SARS-CoV-2 spike (Cat: 848561), positive control Hs-PPIB (Cat: 313901), or negative control DapB (Cat: 310043) were hybridized followed by proprietary assay signal amplification and detection. Tissues were counterstained with Gill’s hematoxylin. An uninfected mouse was used as a negative control and stained in parallel. Tissues were visualized using an Olympus BX60 microscope and imaged with a Nikon (Model #) camera.

### Immunocytochemical staining for SARS-CoV-2

iPSC-derived neurons on glass-bottomed 4-well chamberslides were gently washed with 1X PBS and fixed for 1hr with 4% paraformaldehyde for removal from BSL-3 facility. Wells were subsequently washed 3 times with 1X PBS, permeabilized for 15min at RT in 0.1% Triton-X in PBS and blocked for 2hrs in a 2% BSA-5% NDS blocking solution. Cells were incubated overnight at 4°C in blocking solution with primary antibodies anti-MAP2 (EnCor Biotech. Cat:NC0388389) and anti-SARS-CoV-2 nucleocapsid (Sino Bio. Cat: 40143-R019). Cells were then washed with 1X PBS and incubated in blocking solution for 2hrs at RT with secondary antibodies, Alexa Fluor 594-conjugated goat anti-chicken and Alexa Fluor 488-conjugated goat anti-rabbit. After staining, cells were imaged for presence of SARS-CoV-2 nucleocapsid staining and quantified using the Revolve D75.

## Acknowledgements

This work was supported by a COVID CRAFT grant from UC Irvine Office of Research, NIH-NINDS R35 NS116835, NIH-NINDS NS041249, National Multiple Sclerosis Society grant CA-1607-25040 and support from the Ray and Tye Noorda Foundation to T.E.L. L.M.T. was supported by NIH-NINDS R35 NS116872 and K.N.G. was supported by NIH-NINDS R01NS083801. G.M.O. was supported by Immunology Research Training Grant 5T32AI060573-15 and C.S-G was supported by the Hereditary Disease Foundation. The authors wish to acknowledge the support of the Chao Family Comprehensive Cancer Center Experimental Tissue Shared Resource supported by the National Cancer Institute of the National Institutes of Health under award number P30CA062203.

